# Spinal circuits encode a map of the trunk to control skin twitches

**DOI:** 10.64898/2026.05.10.724081

**Authors:** Rémi Ronzano, Ka Yee Chan, Manon Papadiamandis, Ugo Momi, M. Gorkem Ozyurt, Marco Beato, Robert M. Brownstone

## Abstract

Reflexive skin twitches are stereotypical behaviours, triggered in most mammals by mechanosensory stimuli. The neural circuits controlling this behaviour are thought to lie in the spinal cord, with motor neurons positioned in the lower cervical spinal cord innervating the cutaneous maximus muscle responsible for the twitches. The spatial matching of the motor output to the location of the sensory stimulus points to selective innervation of particular cutaneous maximus motor neurons by specific sensory-responsive circuits organised in a spatially-dependent manner. Using mouse genetics and viral tracing, we observed that a subset of dorso-lateral dI3 neurons forms ascending projections to motor neurons mediating skin twitches. Their projections are somatotopically organized to map a two dimensional space onto specific sub-compartments of the cutaneous maximus motor pool. Furthermore, direct optogenetic stimulation of thoraco-lumbar dI3 ascending projections in the cervical spinal cord induces skin twitches. Together, we demonstrate the circuit basis of a spinal sensory-motor representation of the dorsolateral trunk that participates in an ethological behaviour shared by most mammals allowing them to reduce the burdens of irritants such as insects.

## Introduction

In the famous poem by the French writer La Fontaine, the gnat triumphs over the lion. Although this was primarily written to warn the royal court against signs of pride, this story holds some ground truth. Insects are a deadly threat to mammals. They have a massive impact on livestock (Boonsaen et al., 2024; Taylor et al., 2012) and a similar pressure on wild mammals (Balashov, 2007). Across the course of evolution, the selective pressure exerted by insects and other skin irritants led to the emergence of stereotypical behaviours in mammals, aimed at reducing the detrimental impacts (Dougherty et al., 1995; Woolley et al., 2018). One largely conserved behaviours is reflexive and allows most mammals including horses, zebras, dogs, hedgehogs, and rats, amongst others, to twitch the skin on their backs and flanks, zones inaccessible to scratching by their limbs (Essig et al., 2013; Hall, 1833; Krogh, J. E., & Denslow, 1979; Petruska et al., 2014; Wilson, 1898; Woolley et al., 2018).

This reflexive response, effected by the cutaneous maximus muscle (called cutaneous trunci or panniculus reflex), is triggered by local sensory stimuli leading to local contractions of the portion of the cutaneous maximus muscle located underneath the stimulated region of skin (Krogh, J. E., & Denslow, 1979; Nixon et al., 1984). The cutaneous maximus muscle is a thin layer of skeletal muscle that covers most of the posterior abdomen and thorax of mammals (Figure 1A, Crouch, 1969). Sensory neurons conveying this information to the spinal cord are organised rostro-caudally and as such convey sensory signals to spinal segments between the rostral thoracic and the first rostral half of the lumbar spinal cord (Petruska et al., 2014). Yet motor neurons innervating the cutaneous maximus are confined to the most caudal segments of the cervical spinal cord (Haase & Hrycyshyn, 1985; Holstege et al., 1987; Krogh, J. E., & Denslow, 1979; Theriault & Diamond, 1988a). Therefore, this reflex requires multisegmental spinal circuits spanning the thoracic and lumbar spinal cord and encoding a spatial representation of the back and flank of mammals.

**Figure 1:**
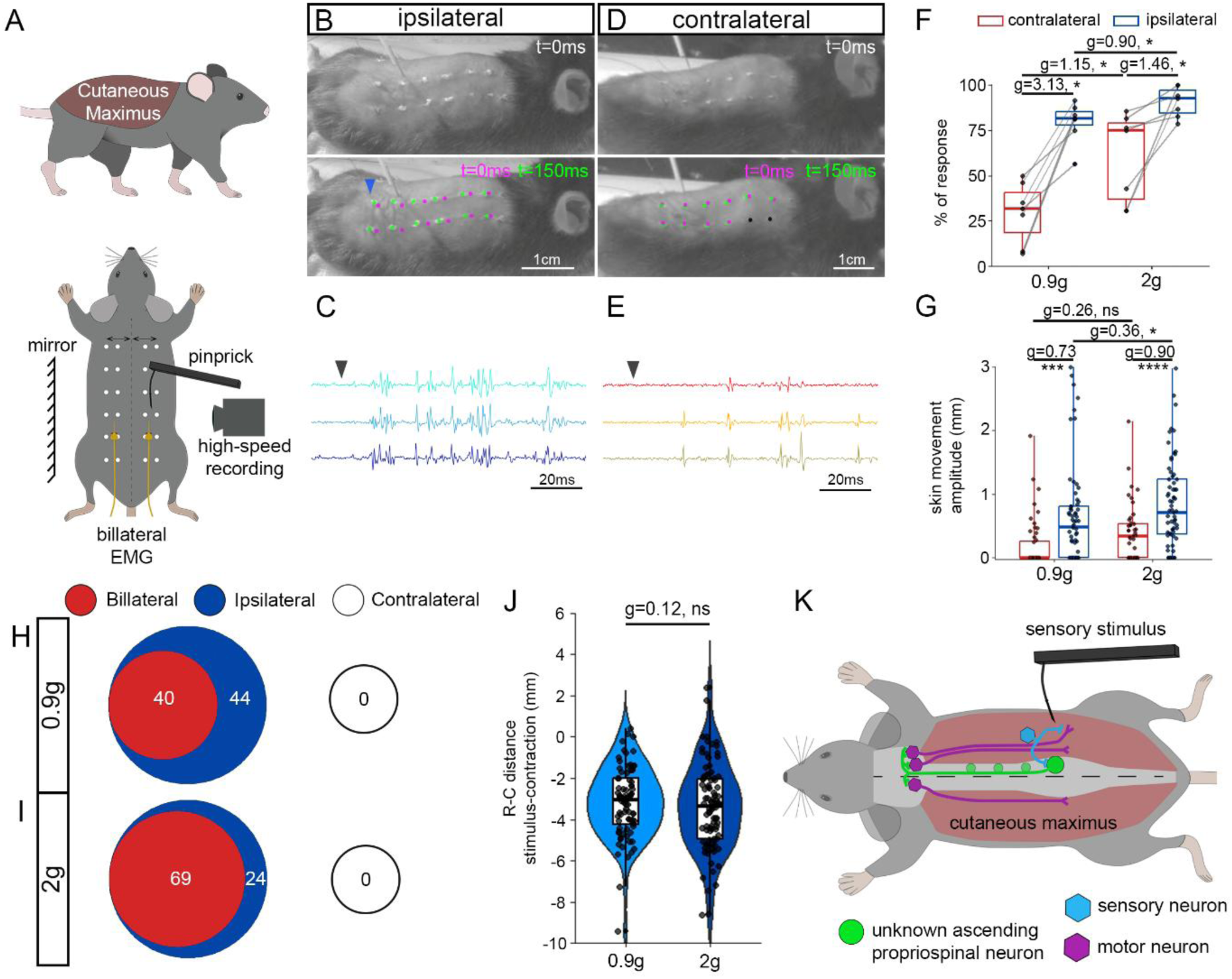
reflexive skin twitches rely on a spatial representation of the torso. (A) Schematics of the position of the cutaneous maximus muscle (top) and of the experimental strategy used to bilaterally record skin movements and cutaneous maximus EMG signals following pinprick stimulation. (B, D) Example of recorded skin movements following stimulation with a 0.9g pinprick, on the side ipsilateral (B) and contralateral (D) to the pinprick. Images at t=0ms are from the first frame with contact of the pin on the skin, while 150ms later the maximum amplitude of the twitch is reached. Magenta and green dots highlight the position of dots placed on the skin to visualize the twitch (magenta for t=0ms, green for t=150ms). Black dots indicate no movement of the dot between the two time points. The blue arrowhead points at the position of initial muscle contraction. (C, E) Differential myomatrix EMG signals recorded from the ipsilateral (C) and (E) contralateral cutaneous maximus from the response shown in (B) and (D), respectively. The black arrowhead indicates the approximate timing of pinprick. (F) Percentage of trial leading to an ipsilateral (blue) versus contralateral (red) skin twitch following stimulation with a pinprick glued to a 2g and 0.9g Von Frey filament. (G) Maximum amplitude of skin movement on the side ipsilateral (blue) and contralateral (red) to the pinprick when animals show a skin twitch reflex. (H-I) Distribution of skin twitch responses bilateral (red), ipsilateral only (blue) and contralateral only (black) following stimulation with a 0.9g (H) and 2g (I) pinprick. (J) Rostro-caudal distance between the sensory stimulation and the location of the initial contraction of the cutaneous maximus following stimulation with a 0.9g and 2g pinprick. When contraction is caudal to the sensory stimulus the distance is set as negative and set as positive if rostral. (K) Based on the behavioural observations, ascending propriospinal neurons (green) need to encode a spatial representation of the torso and convey sensory signals (sensory neuron, blue) to the cutaneous maximus motor neurons (magenta) innervating the part of the muscle located were the sensory stimulus occurred. An additional contralateral pathway, never activated independently from the ipsilateral circuit, recruits the contralateral motor neurons innervating the medial part of the muscle. (F) Individual values plotted from n=7 mice, pairwise Wilcoxon test with Benjamini-Hochberg correction for multiple comparison and bootstrapped Hedge’s G coefficients. (G) Hierarchically bootstrapped Hedge’s G coefficients and Kruskal-Wallis test followed by Dunn’s multiple comparisons tests. n = 70 (2g, ipsilateral), 43 (2g, contralateral), 64 (0.9g, ipsilateral) and 52 (0.9g, contralateral) trials in a total of N=8 mice. (H-I) Response from n=8 mice. (J) n = 92 (2g), and 80 (0.9g) responses in a total of N=5 mice. Non-hierarchically bootstrapped Hedge’s G coefficients and two-tailed t-test.

Sensory stimuli perceived by individuals in their environment are encoded in the central nervous system in topographic maps (Kaas, 1997). The somatotopic organisation of circuits encoding mechanosensory signals has been the focus of many studies from the description of the sensory homunculus (Penfield & Boldrey, 1937) to more recent studies investigating how nervous systems integrate a myriad of spatially discrete sensory signals from the periphery to the brain by somatotopically aligned columns (Chirila et al., 2022; Kuehn et al., 2019; Li et al., 2011; Turecek et al., 2022). Yet, mechanosensory signals are also processed within spinal cord circuits where they recruit topographically organised modules responsible for tuning ongoing motor programs and triggering reflexive movements (Bui et al., 2013; Drew & Rossignol, 1987; Duysens & Pearson, 1976; Gatto et al., 2021; Ozyurt et al., 2025; Quevedo et al., 2005). Most reflexes engage limbs, with sensory stimuli leading to the recruitment of motor neurons controlling discrete points, i.e. joints. In sharp contrast, reflexive skin twitches are triggered by the activation of a restricted part of the cutaneous maximus, which requires a topographical alignment of sensory signals with the activation of motor outputs. Here we sought to define the spinal circuits that are responsible for this stereotypical behaviour and asked how these circuits encode a two dimensional representation of the dorso-lateral trunk. We previously showed that spinal dI3 neurons are mediating cutaneous reflexes such as grasping reflexes (Bui et al., 2013) and stumbling corrections (Ozyurt et al., 2025). This cardinal class of spinal neurons arise from the third dorsal spinal progenitor domain of the notochord, is characterised by expression of the transcription factor *Isl1* (Helms & Johnson, 2003), and is likely composed of several functional subpopulations (Ozyurt et al., 2025). Furthermore, dI3 neurons express classic markers of projection neurons (Osseward et al., 2021), and were suggested to form ascending lumbo-cervical projections that can trigger motor activity in neonates (Nasiri et al., 2025). Here, using mouse genetics, circuit mapping, and behavioural approaches to dissect the neural basis of reflexive skin twitches, we show that a dorso-lateral subpopulation of dI3 neurons form ascending projections to cutaneous maximus motor neurons to evoke reflexive skin twitches. Furthermore, their ascending projections are somatotopically organised along the medio-lateral and rostro-caudal axes and form synapses with cervical motor neurons innervating the cutaneous maximus muscle. This somatotopy encodes a two dimensional spatial map of the thorax and abdomen, providing the circuit foundation for the control of reflexive protective skin twitches.

## Results

### Reflexive skin twitches rely on a spatial representation of the torso

Reflexive skin twitches are one of the main stereotypical behaviours observed in mammals to reduce the burden of skin irritants like insects. To characterize this behaviour in mice, we first applied noxious stimuli (pinprick) on the dorsal and lateral parts of the torso of mice under light sedation. These stimuli were applied using pinpricks of different stiffnesses, while recording movements of the skin and EMG activity of the cutaneous maximus muscles bilaterally (Figure 1A). We observed that these stimuli resulted in activation of the cutaneous maximus muscle on the side of the stimulation, recorded as a rapid movement of the skin (Figure 1B-C; Supplementary figure 1A-B; Supplementary video 1-2). The probability of triggering a skin twitch and the amplitude of the twitch scaled with the force applied: a 2g versus 0.9g pinprick resulted in a higher proportion of responses, increased activity of the cutaneous maximus, and larger skin movements on the side ipsilateral to the stimulation (Figure 1C, F, G; Supplementary figure 1B). Furthermore, as described previously in other species (Blight et al., 1990; Muguet-Chanoit et al., 2011; Theriault & Diamond, 1988b; White et al., 2019), in some instances the sensory stimulation triggered an active movement of the skin on the contralateral side; this response was always smaller compared to the ipsilateral response (Figure 1D-G; Supplementary figure 1C-D, Supplementary video 1-2). The likelihood of contralateral skin twitches also scaled with the force of the pinprick (Figure 1F-G), and in the vast majority of cases led to more pronounced movement of the medial compared with the lateral skin (Supplementary figure 1I). Importantly, these contralateral responses were never seen independently of an ipsilateral twitch. That is, the skin twitch reflex was either ipsilateral or bilateral but never solely contralateral (Figure 1H-I).

To further test whether the reflexive skin twitches relied on a spatial representation of the dermatome overlying the cutaneous maximus muscle, we then measured the rostro-caudal distance between the location of the stimulus and the initial contraction of the cutaneous maximus. The initial contraction could be visualized as the location where the skin on each side of the contraction started getting closer together even in trials in which the recruitment of the muscle spanned a large proportion of the body length (Figure 1B; Supplementary figure 1A). We found that the rostro-caudal distance between sensory stimulation and motor contraction was about 3 mm on average, which represented ∼4.5% of the animal body length with no difference between the 0.9g and 2g pinprick (body length measured from the base of the neck to the base of the tail). Furthermore, the motor contraction site was consistently caudal to the sensory stimulus and was independent of the stimulus strength (3.4 ± 2.1 mm and 3.2 ± 1.7 mm for pinprick 2g and 0.9g respectively, Figure 1J; Supplementary figure 1J).

In some mammals, skin twitches can be triggered by non-noxious sensory stimuli as well. To determine whether this is the case in mice, we used low resistance Von Frey filaments of 0.9g and 2g that are known to primarily activate low-threshold, rather than high threshold, mechanoreceptors (Lezgiyeva et al., 2025; Qi et al., 2024). We observed that Von Frey filaments can induce skin twitches (Supplementary figure 1E-H, Supplementary video 3-4), with the likelihood of the twitch scaling with the mechanical force, similarly to what was observed using pinpricks (Supplementary figure 1K).

Altogether, this shows that noxious and non-noxious stimuli trigger focal skin twitches in mice. Spinal circuits controlling this reflex encode a rostro-caudal representation of the sensory stimuli and are able to recruit specific cutaneous maximus motor neurons in the caudal cervical cord that innervate the location were the sensory stimulus originated (with a slight caudal shift). The contralateral response, also shows that the spinal circuits involved can induce bilateral activation by recruiting cutaneous maximus motor neurons and more specifically those innervating the medial portion of the muscles, thereby spatially encoding the medio-lateral axis of the torso (Figure 1K).

### dI3 neurons form lumbo-cervical projections innervating cutaneous maximus motor neurons

It was first shown in cats that direct lumbo-cervical propriospinal ascending projections to lamina IX seemed to be targeted to a single group of neurons in the caudal cervical cord (Giovanelli Barilari & Kuypers, 1969; Matsushita & Ueyama, 1973; Sterling & Kuypers, 1968), later identified as the cutaneous maximus motor pool (Holstege et al., 1987). Later on, these fibres were shown to largely originate from excitatory propriospinal neurons (Ruder et al., 2016). We reasoned that dI3 neurons express projection neuron markers (Osseward et al., 2021), generate lumbo-cervical motor signals (Nasiri et al., 2025), and mediate disynaptic cutaneous reflexes including grasping reflexes and stumbling reactions (Bui et al., 2013; Ozyurt et al., 2025). Therefore, they are good candidates to form these specific ascending propriospinal projections and be the neuronal substrate responsible for conveying sensory information to cutaneous maximus motor neurons.

To test this hypothesis, we first used bilateral lumbar AAV injections in Isl1-cre mice to induce cre-dependent expression of membrane-tagged YFP to trace the projections of lumbar dI3 neurons, mapping their axons throughout the cervical spinal cord (Figure 2A; Supplementary figure 2A-D). We observed that lumbar dI3 neurons form sparse ascending projections distributed throughout the ventral horn and around the central canal of cervical segments, with dense projections confined to the most ventral part of lamina IX in cervical segments C7 and C8 (Figure 2B-E; Supplementary figure 2E-I). To more specifically visualize pre-synaptic terminals of dI3 lumbo-cervical projections, we used a similar strategy but with a viral construct inducing the expression of a GFP targeted to presynaptic compartments (Figure 2F). Doing so, we first confirmed that the GFP puncta visualized were specific to presynaptic terminals, with the vast majority colocalized with vGlut2 expression (Supplementary figure 2J-K). We then proceeded to map the terminals of lumbar dI3 in cervical spinal segments. Similarly to what we observed with axonal projections, lumbar dI3 terminals were distributed sparsely in the ventral horn and around the central canal of all segments from C3 to the first thoracic segment T1 (Figure 2H-J; Supplementary figure 2L-P). The number of terminals from lumbar dI3 neurons was particularly prominent in segments C7 and C8 due to the dense projections to the ventral lamina IX of these two segments, a pattern consistently observed in every sample (Figure 2H-J). Importantly, these dense projections were segregated to the region of lamina IX that is not reached by proprioceptive axons in these two segments (Figure 2K), previously described as the location of cutaneous maximus motor neurons in mice (Pecho-Vrieseling et al., 2009; Vrieseling & Arber, 2006).

**Figure 2:**
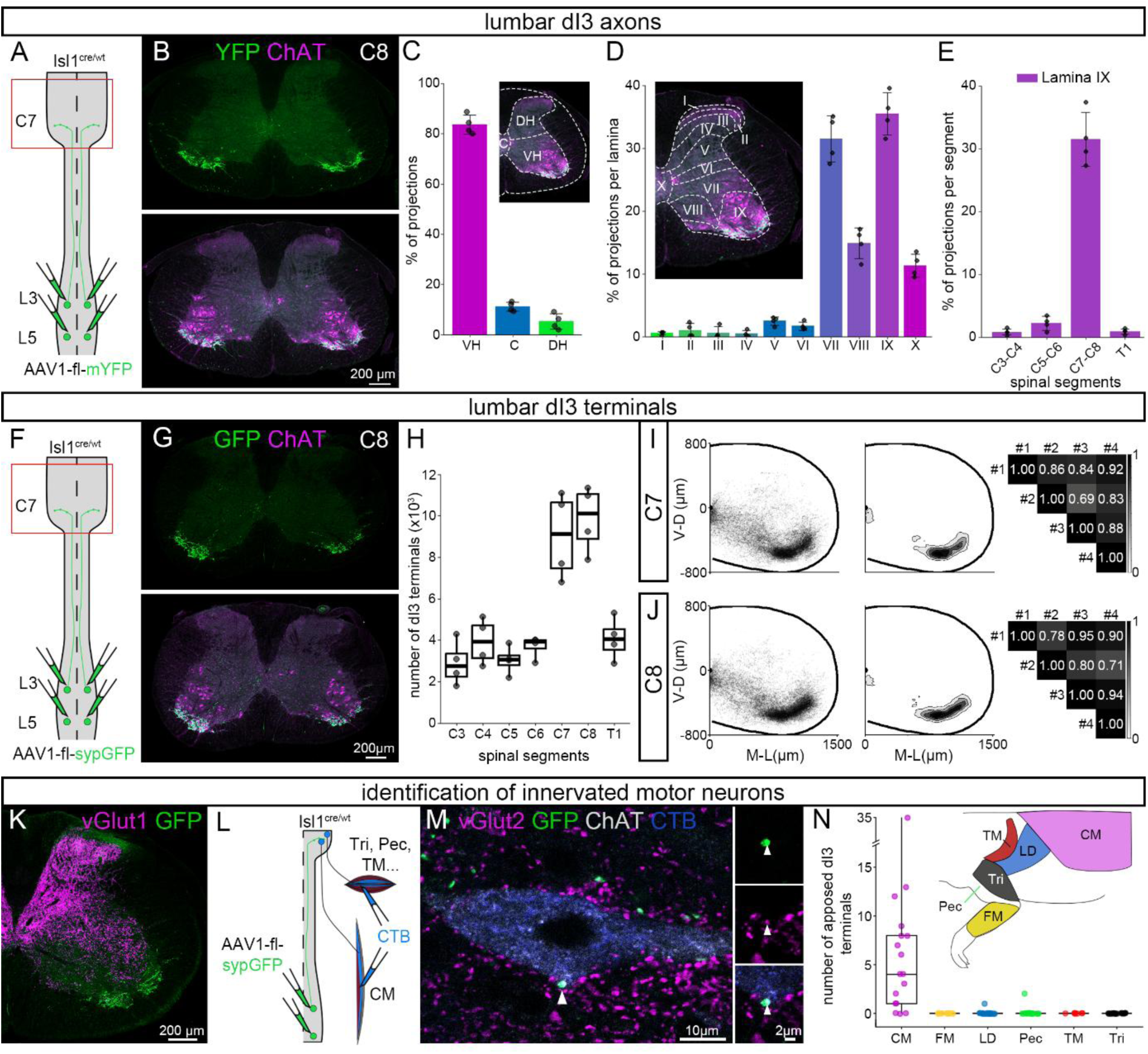
lumbar dI3 neurons form long ascending projections to cutaneous maximus motor neurons. (A) Experimental strategy used to map axonal projections of lumbar dI3 neurons in cervical segments of the spinal cord. (B) Orthogonal projection showing lumbar dI3 axons (YFP, green) and cholinergic neurons (ChAT, magenta) in cervical segment C8. (C) Percentage of lumbar dI3 neurons axonal projections in cervical segments C3 to T1, between ventral horn (VH), central canal area (C) and dorsal horn (DH). (D) Percentage of lumbar dI3 axonal projections in cervical segments C3 to T1, across spinal cord laminae. (C-D) Images insets show plotted subregions on a transverse section. (E) Distribution of lumbar dI3 axonal projections along lamina IX of segments C3 to T1. (F) Experimental strategy used to map lumbar dI3 presynaptic terminals in cervical segments of the spinal cord. (G) Representative orthogonal projection of a transverse section from spinal segment C8 showing lumbar dI3 presynaptic terminals (in green) and cholinergic neurons (in magenta). (H) Number of dI3 terminals (in thousands) from segments C3 to T1 mapped on every other slices. (I-J) Spatial distribution, spatial density and spatial correlation analysis of lumbar dI3 terminals in cervical segments C7 (I) and C8 (J) on an idealized cervical transverse section. (K) Representative coronal slices from cervical segment C8 showing the spatial segregation between lumbar dI3 neurons ascending projections (green) and proprioceptive sensory afferences (ventral vglut1, magenta). (L) Experimental strategy used to visualize presynaptic terminals from lumbar dI3 neurons on identified motor neurons. (M) Single optical slice showing a glutamatergic (vGlut2, magenta) presynaptic terminal from a lumbar dI3 neuron (GFP, green) onto a motor neuron (ChAT, grey) innervating the cutaneous maximus (CTB, blue). The white arrowhead highlights a dI3 bouton apposed to the cutaneous maximus motor neuron, shown at higher magnification on the right side of the panel. (N) Number of lumbar dI3 neuron terminals apposed to identified motor neurons. Schematic shows the position of the targeted muscles; CM: cutaneous maximus, FM: forearm muscles, LD: latissimus dorsi, Pec: pectoralis, TM: teres major, Tri: triceps. (C-E, H) Data shown are from n=4 animals, with individual values plotted. (I-J) Spatial distributions and densities are pooled from tracing in n=4 animals. (N) Individual values correspond to motor neuron, for each muscle with n the number of motor neurons and N the number of mice: CM n=17 in N=3, FM n=17 in N=2, LD n=24 in N=2, Pec n=19 in N=3, TM n=7 in N=2, Tri n=21 in N=2.

To visualize whether lumbo-cervical dI3 projections could directly innervate cutaneous maximus motor neurons, we combined the anterograde labelling of dI3 neuron terminals with retrograde tracing of cutaneous maximus motor neurons (Figure 2L). We observed that the vast majority of cutaneous maximus motor neurons labelled had dI3 terminals apposed to their somata (14/17 in n=3 animals, Figure 2M-N). In contrast, the vast majority of retrogradely labelled neighbouring motor neurons innervating various forelimb muscles showed no dI3 terminals apposed to their somata 86/88 (Figure 2N). Thus, the lumbo-cervical projections of dI3 neurons form an anatomical substrate that could mediate the activation of the cutaneous maximus muscle and thus mediate reflexive skin twitches.

### Ascending projections are formed by a subset of dorso-lateral dI3 neurons

The dI3 cardinal class forms a large population of spinal neurons likely comprising several subpopulations (Ozyurt et al., 2025). To determine which dI3 neurons form ascending projections to cutaneous maximus motor neurons, we injected AAVrg-GFP into the C7 and C8 ventral horns in mice expressing tdTomato in dI3 neurons (Figure 3A). Following the injection, we mapped the distribution of all dI3 neurons (tdTomato^ON^) in the lumbar spinal cord and the ones projecting to the injected site (GFP^ON^tdTomato^ON^). We observed that GFP^ON^tdTomato^ON^ dI3 neurons were situated in the dorso-lateral subpopulation of dI3 neurons (Figure 3E-G), close to the site where Aδ and C-fibers labelled from the dorsal cutaneous nerve, thought to mediate skin twitches, terminate (Lee et al., 2017). Furthermore, dorso-lateral ascending dI3 neurons were absent from segments L5 to L6, very sparse in L4 and consistently distributed from L3 to L1 (Supplementary figure 3A-B). This pattern matches the caudal limit of the cutaneous maximus reflex since it cannot be reliably triggered in dermatomal segments caudal to L4 (rat – Petruska et al., 2014).

**Figure 3:**
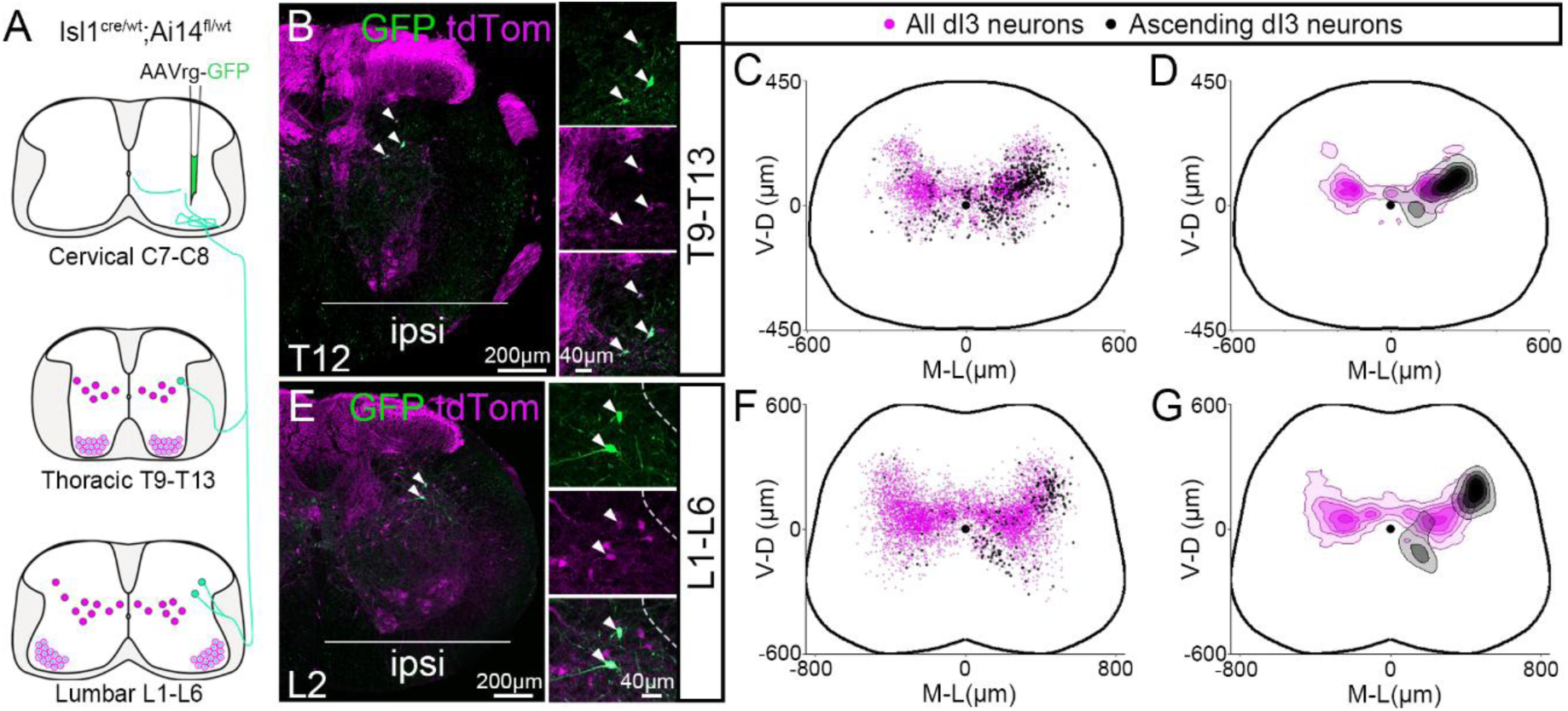
Long ascending projections are formed by a subset of dorso-lateral dI3 neurons. (A) Experimental strategy used to map across thoraco-lumbar segments the global distribution of dI3 neurons (tdTomato^ON^) and the distribution of dI3 neurons projecting to the ventral horn of C7-C8 segments (GFP^ON^tdTomato^ON^). (B, E) Orthogonal projections of ipsilateral hemislices showing GFP^ON^tdTomato^ON^ dI3 neurons (GFP green, tdTom magenta) from thoracic segment T12 (B) and lumbar segment L2 (E). GFP^ON^tdTomato^ON^ dI3 neurons are shown at higher magnification on the right side of the panels. (C-G) Spatial distributions (C, F) and spatial densities (D, G) of all dI3 neurons (magenta) and dI3 neurons projecting to the ventral horn of segment C7-C8 (black). Maps are overlapped distributions from segments T9 to T13 (C-D) and L1 to L6 (F-G) on idealized transverse thoracic and lumbar sections respectively. Results pooled from n=4 mice, mapped on every other sections for GFP^ON^tdTomato^ON^ dI3 neurons while the global distribution of dI3 neurons (tdTomato^ON^) was mapped on one section every four.

We next sought to determine whether ascending dI3 neurons are distributed along thoracic spinal cord by mapping retrogradely traced dI3 neurons in caudal (T9 to T13) and rostral (T3 to T6) segments. Ascending dI3 neurons were present throughout the thoracic spinal cord. Similarly to lumbo-cervical projecting dI3 neurons, thoracic GFP^ON^tdTomato^ON^ dI3 neurons in segments T9 to T13 were distributed in the dorso-lateral region of the dI3 population (Figure 3B-D; Supplementary figure 3A-B). In rostral thoracic segments, this dorso-lateral population of ascending dI3 neurons was mixed with a population located ventro-medially, which may represent short ascending dI3 neurons involved in other circuits (Supplementary figure 3C-F).

Altogether, our results show that ascending dI3 neurons projecting to the location of the cutaneous maximus motor pool form a column of dorso-lateral neurons from rostral thoracic segments to the rostral half of the lumbar cord, a distribution that fits with the segmental organisation of the skin twitch reflex.

### Ascending projections of dI3 neurons are somatotopically organised to encode a spatial map of the trunk

The vast majority of retrogradely labelled dI3 neurons were located ipsilateral to the side of injection. Nevertheless, a few dI3 neurons were also found contralaterally (Figure 3C, F; Supplementary figure 3E). We reasoned that ascending dI3 neurons projecting contralaterally could therefore mediate the contralateral skin twitches observed behaviourally. To test this hypothesis, we mapped presynaptic terminals from unilaterally transduced dI3 neurons between T12 and L2 (Figure 4A) and found that they densely innervate the ipsilateral cutaneous maximus motor pool and also form contralateral, but less abundant, projections segregated to the same domain (Figure 4B, D). The same pattern was obtained when unilaterally targeting dI3 neurons located between thoracic segments T9 and T11 with a similar spatial distribution in the transverse plane (Figure 4A, C-F; Supplementary figure 4A) and about 15% of the cervical terminals of dI3 neurons located contralaterally (Figure 4G). Mapping every other sections in C7 and C8 segments, we found ∼20 000 dI3 terminals per animal, that taking into account the number of dI3 neurons transduced corresponded to a minimum of 180 terminals per dI3 neuron (Supplementary figure 4B-C). To confirm that these projections bilaterally innervate cutaneous maximus motor neurons, we retrogradely labelled them bilaterally following an unilateral tracing of presynaptic terminals from thoraco-lumbar dI3 neurons (Supplementary figure 4D). We observed that ipsilateral cutaneous maximus motor neurons frequently (25/27) showed multiple terminals of dI3 neurons apposed to their soma (Supplementary figure 4E, G-H). Additionally, although terminals apposed to contralateral cutaneous maximus motor neurons were less frequent, about 40% showed dI3 terminals apposed to them (8/21, Supplementary figure 4F-H). Therefore, the anatomical evidence points to ascending dI3 neuron projections being able to recruit cutaneous maximus motor neurons and trigger skin twitches on both sides of the body.

**Figure 4:**
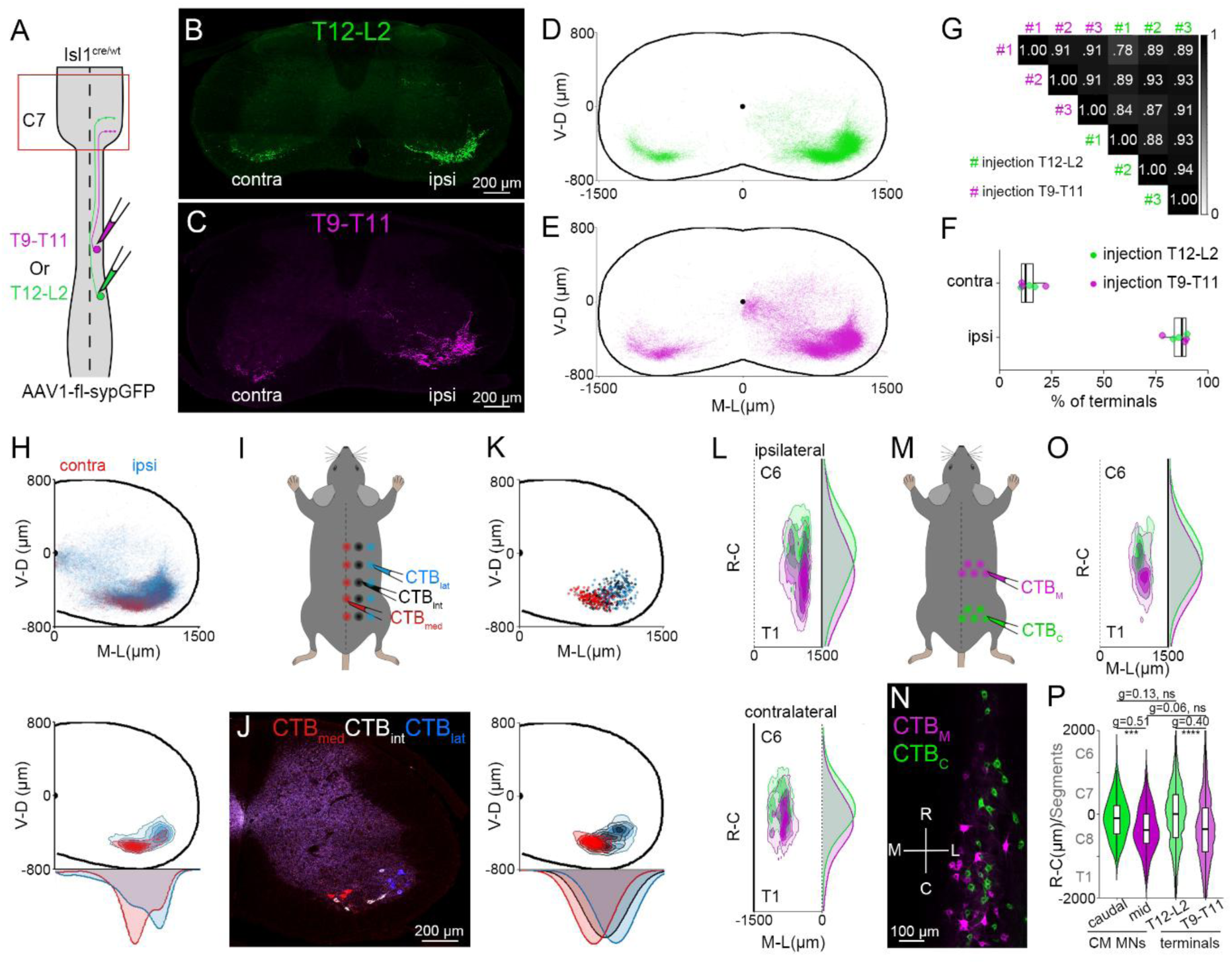
Cutaneous maximus motor neurons and ascending dI3 projections are somatotopically organized along the medio-lateral and rostro-caudal axis. (A) Experimental strategy used to map presynaptic terminals of thoraco-lumbar dI3 neurons in cervical segments of the spinal cord. (B-C) Orthogonal projection showing presynaptic terminals (GFP) of dI3 neurons from segments T12 to L2 (B, green) and segments T9 to T11 (C, magenta) on cervical transverse sections representative of segments C7 and C8. (D-E) Spatial distributions of GFP presynaptic terminals from dI3 neurons in T12 to L2 (D) and T9 to T11 (E) mapped from segments C6 to T1 on an idealized cervical transverse section. (F) Spatial correlation analysis of dI3 presynaptic terminals on the transverse plane in segments C6 to T1 from dI3 somata in segments T12 to L2 (green) and T9 to T11 (magenta). (G) Percentage of dI3 presynaptic boutons on the ipsilateral versus contralateral side of the injection analysed from segments C6 to T1. Individual values are plotted as green dots for tracing from T12 to L2 dI3 neurons and in magenta for dI3 neurons in segments T9 to T11. (H) Spatial distribution (top) and spatial density (bottom) of dI3 neurons terminals mapped from segments C6 to T1, with an overlap on an idealized transverse hemisection of ipsilateral (blue) and contralateral (red) distributions. (I) Schematic of the experimental strategy used to retrogradely label motor neurons innervating the medial (red), intermediate (black) and lateral (blue) part of the cutaneous maximus. (J) Orthogonal projection of a representative cervical transverse section showing motor neurons innervating the medial (red), the intermediate (white) and the lateral (blue) part of the cutaneous maximus muscle. (K) Spatial distribution (top) and density (bottom) of motor neurons innervating the medial (red), intermediate (black) and lateral (blue) part of the cutaneous maximus on an idealized cervical transverse section. (L) Spatial density of presynaptic terminals from dI3 neurons of T9 to T11 segments (magenta) and T12 to L2 segments (green) along the rostro-caudal axis on an idealized horizontal section from segments C6 to T1. Densities are plotted independently for terminals on the ipsilateral (top) and contralateral (bottom) side to the injection. (M) Schematic of the experimental strategy used to retrogradely label cutaneous maximus motor neurons innervating the middle (magenta) and caudal (green) part of the muscle. (N) Orthogonal projection of a horizontal section showing motor neurons innervating the middle (magenta, CTB_M_) and caudal (green, CTB_C_) part of the cutaneous maximus. M-L indicates the medio-lateral axis while R-C gives the orientation along the rostro-caudal axis. (O) Spatial density of motor neurons innervating the caudal (green) and middle (magenta) part of the cutaneous maximus along the rostro-caudal axis on an idealized horizontal section from segments C6 to T1. (P) Plot showing the rostro-caudal distributions of presynaptic terminals from dI3 neurons in T9-T11 (magenta) and T12-L2 (green) and motor neurons of the cutaneous maximus (CM MNs) innervating the middle (magenta) and caudal (green) part of the muscle. (D-E, K-L, O) Spatial distributions and densities are pooled from tracing in n=3 mice for dI3 terminals and n=4 mice for cutaneous maximus motor neurons. (G) Data shown are from a total of n=6 mice with individual values colour coded by type of injection. (H) Spatial distribution is from n=6 mice with n=3 mice for each type of injection (T9-T11 and T12-L2). (P) Distributions from n=4 and n=3 mice for motor neurons somata and dI3 neurons terminals respectively. Non-hierarchically bootstrapped Hedge’s G coefficient and Kruskal-Wallis test followed by Dunn’s multiple comparisons test performed on somata/terminals.

How do skin twitches induce a local contraction of the cutaneous maximus muscle that depend on the location of the sensory stimulus medio-laterally and rostro-caudally? To determine the circuits architecture encoding spatial information, we first focused on contralateral projections of dI3 neurons. Behaviourally, contralateral responses of the medial skin show a higher amplitude of movement than that of the lateral skin (Supplementary figure 1I). We reasoned that a medio-lateral organisation of ascending dI3 projections that would mirror the motor neuron distribution could form the substrate to induce skin twitches at specific locations along the medio-lateral axis. To test this hypothesis, we overlapped the distribution of ipsilateral and contralateral ascending dI3 terminals in the caudal cervical cord and compared their density along the medio-lateral axis. This unravelled a contralateral distribution located medially compared to the ipsilateral distribution that spanned the whole medio-lateral domain of the cutaneous maximus motor pool (Figure 4H). Next, to validate previous retrograde tracing from the different branches of the lateral thoracic nerve (Pan et al., 2012), we retrogradely traced cutaneous maximus motor neurons from the lateral, intermediate and medial part of the muscle (Figure 4I). Supporting previous observations (Pan et al., 2012), these tracings showed a somatotopic organisation of cutaneous maximus motor neurons along the medio-lateral axis (Figure 4J-K). Therefore, the somatotopic organisation of motor neurons together with the medio-lateral distribution of ascending dI3 projections spatially encode information along the medio-lateral axis of the animal.

Yet skin twitches also match the position of the sensory stimulation along the rostro-caudal axis (Figure 1J; Supplementary figure 1J). Therefore, we sought to determine whether ascending dI3 projections could similarly encode space by mirroring a potential somatotopic rostro-caudal organisation of cutaneous maximus motor neurons. To test this hypothesis, we mapped terminals from dI3 neurons (T9-T11 and T12-L2) in the cutaneous maximus pool. We observed that the two distributions span a large part of C7 and C8 segments, which could explain how high amplitude skin twitches span a large part of the rostro-caudal extent of the muscle. However, these two distributions on both the ipsilateral and contralateral sides were shifted, with T9-T11 dI3 neurons projecting to more caudal cervical segments than T12-L2 dI3 neurons (Figure 4L; Supplementary figure 4I). This somatotopic organisation of ascending dI3 neurons terminals along the rostro-caudal axis was further confirmed by comparing individual animals injected at different thoracolumbar level. This analysis revealed a correlation between the position of transduced thoraco-lumbar dI3 somata and the rostro-caudal distribution of their terminals in the cervical spinal cord, with dI3 terminal position that shifts linearly with the position of their somata (Supplementary figure 4J-K). To assess if this rostro-caudal somatotopy mirrored the spatial distribution of motor neurons innervating the caudal versus rostral part of the cutaneous maximus muscle, we retrogradely traced them from the part of the cutaneous maximus muscle underlying the dermatome of spinal segments T9-11 and T12-L2 (Figure 4M). Mapping retrogradely traced motor neurons, we observed a rostro-caudal shift in their distribution (Figure 4N-O; Supplementary figure 4L), with rostro-caudal distributions that matched the ones of ascending dI3 neuron terminals (Figure 4P).

Altogether, these results show a two dimensional somatotopic organisation of ascending dI3 neurons along the medio-lateral and the rostro-caudal axis, mirroring the organisation of cutaneous maximus motor neurons. This mirrored somatotopic organisation, provides the anatomical substrate for the encoding of a two-dimensional spatial representation of the dermatome overlying the cutaneous maximus muscle, which is necessary for the control of reflexive skin twitches.

### Optogenetic stimulation of dI3 ascending projections induce a skin twitch reflex

Based on our anatomical circuit tracings, ascending projections of dI3 neurons could trigger reflexive skin twitches that recapitulate all the features observed at the behavioural level. Therefore, we next tested if specific activation of ascending dI3 neuron projections to the cervical cord is sufficient to trigger skin twitches. To do so, we targeted thoraco-lumbar dI3 neurons with a Cre-dependant AAV construct inducing the expression of the excitatory opsin CoChR (Antinucci et al., 2020; Klapoetke et al., 2014). A month after injection, an optic fibre was positioned just dorsal to the cutaneous maximus motor pool, and optogenetic stimuli were delivered while recording skin movement (Figure 5A). Skin twitches were triggered reliably in most animals (5/6), with the amplitude of the response scaling with the duration of the pulse (Figure 5B-C, Supplementary video 5). Furthermore, the proportion of pulses triggering skin twitches also scaled with the duration of the pulse, with a skin twitch always triggered by 20ms pulses in 4/5 animals (Figure 5D). Lastly, to verify that the skin twitches were indeed evoked by the specific activation of ascending dI3 axons, we performed the same experiments following injections in Cre negative animals. In these animals, we could not induce any skin twitch response even with longer pulses of light, which confirmed that light induced skin twitches in Isl1^cre/wt^ animals were indeed induced by the specific activation of ascending dI3 neuron projections (Figure 5D).

**Figure 5:**
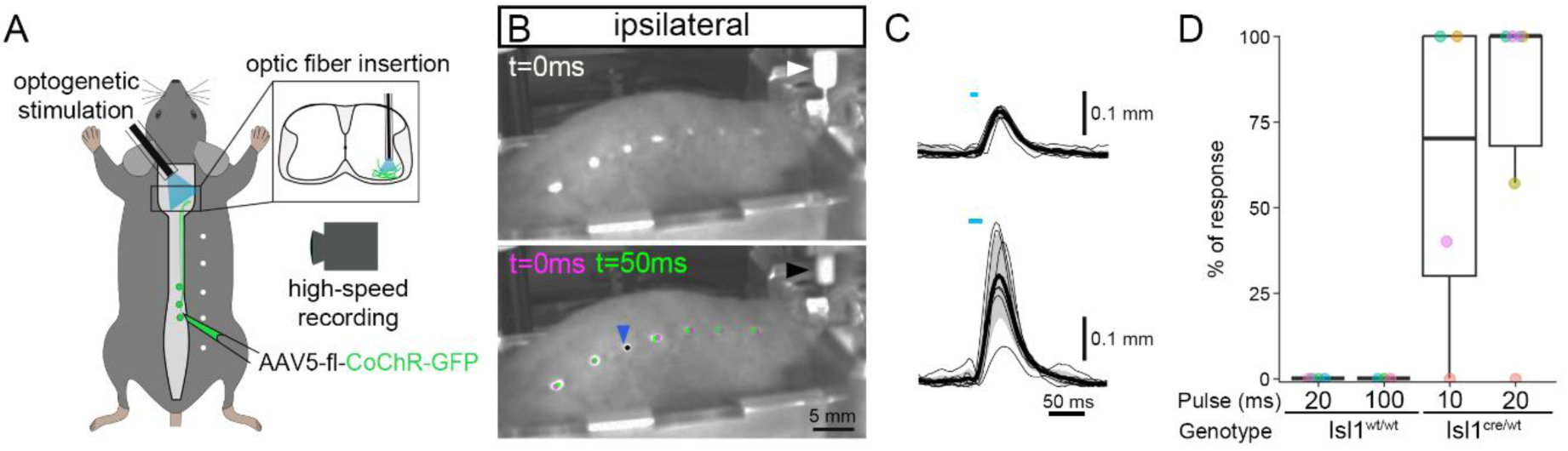
Optogenetic activation of dI3 neurons ascending axons trigger skin twitches. (A) Schematics of the experimental strategy used to optogenetically stimulate ascending projections of dI3 neurons in the caudal cervical cord. The AAV5-fl-CoChR-GFP was injected 4 to 5 weeks before the optic fiber was implanted and the optogenetic stimulations performed. (B) Example of recorded skin movements following an ipsilateral 20ms pulse stimulation. Image at t=0ms are from the first frame with light on (white arrowhead), while 50ms later the maximum amplitude of the twitch is reached. Magenta (t=0ms) and green (t=50ms) dots highlight the position of dots placed on the skin to visualize the twitch. The black dot indicates no movement of the dot between the two time points. The blue arrowhead points the position of initial muscle contraction. (C) Trajectory of the dot with the maximum amplitude along the rostro-caudal axis when optogenetic stimulations are triggered, following a 10ms (top) and a 20ms (bottom) light pulse. The thin lines represent the response to each stimulation in the animal shown in (B), and thick lines are the means of the trajectories. The blue bars show the timing of the light. (D) Percentage of light pulses triggering a skin twitch, colour coded per animal. Pulse indicates the pulse length in milliseconds, and the genotype whether the opsin is expressed (Isl1^cre/wt^) or not (Isl1^wt/wt^). N=4 animals for Isl1^wt/wt^ and Isl1^cre/wt^ for 10ms pulses and N=6 animals for 20ms pulses in Isl1^cre/wt^.

Collectively, our results demonstrate that ascending projections of dI3 neurons can encode spatially restricted sensory signals and trigger reflexive skin twitches that match sensory stimuli location using a two dimensional somatotopic organisation.

## Discussion

Reflexive skin twitches are an ethologically conserved behaviour used by non-primate mammals to reduce the burden of skin irritants like insects on the dorsal and lateral surfaces of their trunk, as these regions are inaccessible to their limbs. Here, we show that dorso-lateral dI3 neurons distributed from the mid-lumbar through thoracic spinal cord form dense ascending projections to cutaneous maximus motor neurons, and that they trigger skin twitches when activated. The spatial organisation of these projections in two dimensions, along the medio-lateral and the rostro-caudal axes, mirrors the somatotopic organisation of the cutaneous maximus motor pools, thus providing the spatial encoding required for effective skin twitches.

Ascending dI3 axons are largely restricted to the region occupied by cutaneous maximus motor neurons with their projections that are organised somatotopically in this spatially confined domain. This topographical organisation raises the question of which developmental signals are at play. Some of the key developmental mechanisms that determine the directionality of long projecting neurons and the position of motor neurons have previously been described (Chédotal, 2019; Demireva et al., 2011; Dewitz et al., 2018; Mendelsohn et al., 2017), but determinants of refined topographical organisation are still unclear. A subset of ascending dI3 neuron axons cross the midline, whereas dI3 neurons have been described as ipsilateral-projecting neurons (Avraham et al., 2010), with the expression of Isl1 supressing the expression of Robo3, important for midline crossing (Masuda et al., 2024; Sabatier et al., 2004). While it is plausible that this small subset of dI3 neurons express Robo3, a Robo3-independent mechanism has been described for lamina IV neurons, with their axons crossing the midline dorsally (Comer et al., 2015; Petkó et al., 2004). Therefore the description of dI3 axons trajectories together with their description of axonal guidance molecules would be helpful to resolve these questions.

Although the twitch reflex is conserved across non-primate mammals, there are some inter-species differences. The rostro-caudal somatotopic organisation of the cutaneous maximus motor pool seems to be inverted compared to the distribution reported in rats where caudal motor neurons innervate the caudal part of the muscle (Theriault & Diamond, 1988a). Another inter-species difference hold in the sensory signals that trigger skin twitches. In rats, they are triggered by noxious stimuli activating Aδ and C-fibers, while a light touch is sufficient in guinea pigs (Blight et al., 1990; Borgens et al., 1987; Nixon et al., 1984; Petruska et al., 2014; White et al., 2019). Our results show that in mice, stimulation of the skin by Von Frey filaments that exclusively trigger activity in low-threshold mechanosensory neurons (Lezgiyeva et al., 2025; Qi et al., 2024) were sufficient to induce skin twitches, even though in our experiments a sensitization may have been facilitated by the presence of the small skin incision used to implant the EMG probe (Duarte et al., 2005). These interspecies differences might be due to differences in the position of dI3 neurons involved, the spatial distribution of mechanosensory terminals, and/or the signalling controlling the specificity of mechanosensory inputs to dI3 neurons. Altogether the circuits for the control of skin twitches emerge as an ideal model to unravel, in future studies, the refined guidance mechanisms organising the topography of the developing spinal cord.

In this study, we have shown that ascending dorso-lateral dI3 neurons trigger reflexive skin twitches. However, future studies aimed at specifically ablating dI3 neurons will be necessary to definitively show whether ascending dI3 neurons form a unique module responsible for skin twitches. Furthermore, it is interesting to note that inhibitory lumbo-cervical projections from an unidentified population have also been showed to terminate in the cutaneous motor pool (Ruder et al., 2016), although sparsely. The role for such a system remains to be clarified, but it could conceivably prevent the reflex in specific conditions.

## Acknowledgement

Myomatrix arrays were supplied by the Center for Advanced Motor BioEngineering and Research (CAMBER) supported by NIH grants U24NS126936 and R01NS109237. We thank Dr. Christopher Black for technical assistance on optogenetic experiments. This work was supported by a Wellcome Trust Early Career Award (225674/Z/22/Z) to R.R. and a Wellcome Trust Discovery Award (227433/Z/23/Z) to R.M.B. and M.B.; R.M.B is supported by Brain Research UK.

## Methods

### Animals

All experiments were performed according to the Animals (Scientific Procedures) Act UK (1986) and certified by the UCL AWERB committee, under project licence number PP2688499. Wild type C57Bl6/J mice, heterozygous *Isl1^Cre/wt^* (Jackson Lab, stock #024242) as is or crossed with homozygous Ai14 (Jackson Lab, stock #007914) were used. All the mice used were from a C57Bl6/J genetic background. All mice were housed with 12 hours of light from 7am to 7pm.

### Tissue collection and immunohistochemistry

The mice were perfused with PBS (0.1 M) followed by PBS 4 % paraformaldehyde (PFA, ThermoFisher, 28908) under terminal ketamine/xylazine anaesthesia (i.p. 80 and 10 mg/kg, respectively). The spinal cords were collected and post-fixed for 2-3 hr, cryoprotected in 30 % sucrose in 0.1M PBS, embedded in PolyFreeze (Polysciences, 25113-1) and sliced coronally (40-50 µm thickness) with a cryostat (Leica, CM3050 S). To quantify the number of dI3 presynaptic boutons onto motor neurons, a step of antigen retrieval was performed prior to the staining protocol using a treatment with citrate buffer 0.3% (ThermoFisher, 036439.A3), pH 6, at 80°C for 3 minutes. Sections were then transferred for a minimum of 30 minutes in PBS 2-X (0.2 M), 0.2-0.3 % Triton 100-X (Sigma, T9284), 10 % donkey normal serum (Sigma, D9663), incubated with primary antibodies for 72 hr at 4°C and with secondary antibodies 24 hr at 4°C in the same blocking solution. For all the other immunohistochemistry, sections were directly transferred in the blocking solution for at least 30 minutes and then incubated with primary antibodies for 32-36 hours at 4°C and with secondary antibodies overnight at 4°C.

The primary antibodies used were: goat anti-choline acetyltransferase (ChAT, 1:100, Millipore, AB144P), rabbit anti-ChAT (1:3750 to 1:7500, Columbia University #CU1574, RRID:AB_2750952), chicken anti-mCherry (1:2000, Abcam, Ab205402), rat anti-RFP (1:1000, Chromotek, clone 5F8), chicken anti-GFP (1:1000, Abcam, Ab13970), rabbit anti-GFP (1:2000, Abcam, Ab290), goat anti-GFP (1:500, Abcam, ab6673), guinea pig anti-GFP (1:750, Synaptic systems, 132 005), goat anti-CTB (1:5000, Nittobo Medical Co, MSFR103230), rabbit anti-CTB (1:400, Abcam, ab34992), rabbit anti-vGlut1 (1:1000, synaptic system, 135 303), guinea pig anti-vGlut1 (1:2000, Millipore, AB5905), and guinea pig anti-vGlut2 (1:2000, Millipore, AB2251-I); and the secondary antibodies: donkey anti-goat Alexa 647 (1:1000, ThermoFisher, A-21447), donkey anti-goat Alexa 488 (1:1000, ThermoFisher, A11055), donkey anti-goat preadsorbed Alexa 405 (1:200, Abcam, Ab175665), donkey anti-guinea pig Alexa 647 (1:1000, Jackson ImmunoResearch, 706-605-148), donkey anti-guinea pig Cy3 (1:500, Millipore, AP193C), donkey anti-guinea pig Alexa 488 (1:500, Jackson ImmunoResearch, 706-545-148), donkey anti-guinea pig DyLight 405 (1:400, Jackson ImmunoResearch, 706-475-148), donkey anti-rabbit Alexa 488 (1:1000, Thermo Fisher, A21206), donkey anti-rabbit Alexa 647 (1:500, Abcam, ab150063), donkey anti-rabbit Alexa 405 (1:500, Abcam, 175649), donkey anti-chicken Cy3 (1:1000, Jackson ImmunoResearch, #703- 165- 155), donkey anti-chicken CF 488A (1:500, Sigma, SAB4600031), donkey anti-rat Alexa 555 (1:500 to 1:1000, Thermo Fisher, A78945). In some experiments, Neurotrace Blue (1:100 to 1:200, ThermoFisher, N21479) was used together with the secondary antibodies. The slides were mounted in Mowiol (Sigma, 81381- 250 G) and coverslipped (VWR, #631- 0147) for imaging.

### Intraspinal injections

Lumbar injections of AAVs were performed injecting P11-P16 animals while retrograde tracing from C7-C8 were performed in 2-4 months old Isl1^cre/wt^;Ai14^fl/wt^ mice under general anaesthesia with isoflurane (5% for induction, 1-1.5% maintenance) with previous induction of analgesia (Buprenorphine, Buprecare, XVD 130). Animals were placed in a custom device for stabilisation of the vertebrae and injected using a pulled and bevelled borosilicate glass pipette (∼30°, Warner instruments, #203-776-0664) with an outer diameter of 30-40µm at the tip, attached to a stereotaxic device (Narishige, SMM-200). AAVs were injected applying multiple successive short pulses using a pneumatic picopump (World Precision Instruments, PV820) at a rate of 100-150nl/min. The glass pipette was then left in place for 1-3 minutes and again 1-3 minutes 50-70µm above the injection sites.

For bilateral injections to trace axonal projections or to visualize presynaptic terminals, 350nl of AAV1-phSyn1-flex-Lyn/EYFP-T2A-synaptophysin:mKate2 (3.2 x 10^11^ vp/mL, VectorBuilder, ID VB180425-1066vcf, expression of mKate2 could not be visualized and therefore only the EYFP signal targeted by the Lyn domain to the membrane was used, abbreviated AAV1-fl-mYFP) or AAV1-hSyn-flex-sypGFP (3.7 x 10^11^ vp/mL, VCF Charité, from construct #153203, a gift from Lynette Lim to Addgene, abbreviated AAV1-fl-sypGFP) respectively were injected per site following a dorsal laminectomy of T13. The injections were performed in the intermediate lamina (380-400µm medio-laterally, 600µm dorso-ventrally) with two sites per side between L3 and L5. The dI3 soma labelled by these injections were quantified in one every other sections from segments C3 to S1, with for AAV1-fl-mYFP injections 1065 ± 300, 8 ± 14 and 4 ± 3 dI3 neurons mYFP^ON^ in the lumbar, sacral and thoracic cord respectively and 1150 ± 199, 2 ± 2 and 9 ± 8 dI3 neurons GFP^ON^ in the lumbar, sacral and thoracic cord respectively following AAV1-fl-sypGFP injection. No positive cells (mYFP^ON^ or GFP^ON^) were found in the cervical part and the ones in the thoracic segments were restricted to the most caudal segments. For unilateral transduction of T12 to L2 dI3 neurons, 150nl of AAV1-hSyn-flex-sypGFP at the same final titer was injected using the laminar spaces between T11-T12 vertebrae (550µm medio-laterally, 400µm dorso-ventrally), no dI3 neurons were transduced outside of T12-L2. For unilateral tracing in spinal segments T9 to T11 dI3 neurons, the same volume was injected in the space between T8 and T9 vertebrae (400µm medio-laterally, 350µm dorso-ventrally). For optogenetic experiments, a full dorsal laminectomy of T12 and a partial laminectomy of T11 or T9 were performed, then 2 sites between T11 and T13 (500µm medio-laterally, 550µm dorso-ventrally) and 2 other sites located below T11 (500µm medio-lateraly, 550µm dorso-ventrally, 4 animals) or T9 (400µm medio-laterally, 350µm dorso-ventrally, 2 animals) were injected with 150nl per site of AAV5-Syn-Flex-CoChR-GFP (8.8 x 10^12^ vp/mL, UNC Viral Vector, lot AV6473) in Isl1^cre/wt^ animals. For control experiments, 4 Cre negative animals (P11-P14) were injected similarly.

Retrograde tracing of dI3 ascending neurons were performed injecting 400-500nl of AAVretro-hSyn-GFP (1.5 x 10^13^ vp/mL, #50465, Addgene) together with CTB-Alexa 647 (0.05%, C34778, ThermoFisher) that was only used to visualize the extent of the site of injections. Injections were performed in a single site (750µm medio-laterally, 1000µm dorso-ventrally), below C7 vertebrae following a dorsal laminectomy.

To control the specificity of Cre dependent AAV constructs, Cre negative animals were injected using the procedure involving the highest volume of injection. At least 2 Cre negative animals per construct were used and not more than 5 fluorescent cells were observed at the sites of injection, while labelling in Isl1^cre/wt^ systematically resulted in the labelling of a large number of cells.

For all experiment involving anatomical analysis mice were perfused three weeks (21-22 days) after the injection of the AAV vectors.

### Intramuscular and intradermic injections

Intramuscular and intradermic injections of CTB were performed 3-10 days before tissue collection in P20 to P40 animals. Animals were anaesthetized using isoflurane inhalation. For intramuscular injections, an incision was made on the skin to visualize the targeted muscle while intradermic injections were made pinching the skin without incision. CTB-biotin (C34779, ThermoFisher), CTB-AlexaFluor-555 (C34776, ThermoFisher), CTB-AlexaFluor-488 (C34775, ThermoFisher) and CTB-AlexaFluor-647 (C34778, ThermoFisher) were used to perform intramuscular (at 0.5%) and intradermic (at 0.20 to 0.25%) injections. Injections were performed using a 5μl Hamilton syringe (model 7652-01) fixed to a manual Narishige micromanipulator (M-3333) and loaded with a bevelled glass (∼30°) pipette of outer diameter 40-50μm. The volume injected and the number of injected sites in each muscle was adapted to the targeted muscle as follow: forearm muscles total of 3μL in 2 sites, triceps total of 4μL in 2 sites, teres major total of 2μL in 2 sites, pectoralis total of 3μL in 3 sites, latissimus dorsi total of 4μL in 3 sites. For all intradermic injections, 0.5μL to 0.7μL were injected per site. For the visualisation of dI3 presynaptic boutons onto cutaneous maximus motor neurons, 5-6 sites per side were injected when tracing from both side of the animals, while 5-9 sites were injected when tracing from only one side. The intradermic injections were performed on the posterior third of the trunk when a concurrent tracing of dI3 in T12 to L5 segments was performed. To map the distribution of cutaneous maximus motor neurons depending on the region of the muscle they innervate, 5 sites were injected for each Alexa Fluor used. For the rostro-caudal map, sites of injections were distributed along the medio lateral axis, with the caudal sites above the sacral and coccygeal vertebrae and the rostral sites above T11 to T13 vertebrae (Figure 4M). For the medio-lateral mapping, for each CTB conjugate, sites were distributed rostro-caudally between the two posterior third of the trunk, with the medial injections on the midline, the intermediate around 0.5 cm lateral and the lateral injections at least 1 cm from the midline.

### Imaging and analysis

All images were obtained using a Zeiss LSM800 confocal microscope. Images of the entire coronal sections were obtained using a x10 air objective (0.3 NA). High magnification images to visualize and quantify presynaptic boutons apposed to motor neurons or colocalization between GFP and vGlut2 were taken using a 63x oil objective (1.4 NA). Tiles were stitched using Zen Blue (ZEN Blue 2.3 software) and analyses were performed using Imaris (Bitplane, version 9.6.1), ImageJ (version 1.54f) softwares and an adapted version of SpinalJ (Fiederling et al., 2021).

Cell and presynaptic terminal mappings were performed respectively manually or semi-automatically using Imaris “Spots” function along spinal segments as specified in the text and on the figures. All distributions were mapped on every other section except for the dI3 global population (tdTomato^ON^) following retrograde tracing that was performed on one section every four. Distribution maps were obtained setting the central canal as (0,0) in the (x,y) Cartesian system with the y-axis set dorso-ventrally, the x-axis medio-laterally and exporting the “Spots” in this Cartesian system. The z position was set for all the structures within the same section depending on the rostro-caudal position of the section in the sample. Positive values were assigned for rostral structures in the z-axis, dorsal in the y-axis and ipsilateral in the x-axis (the x-axis direction was randomly assigned following bilateral injections). The normalization of x and y coordinates was performed independently for each quadrant using the following reference points: the x dimension was normalized to the outer edge of the white matter at the level of the central canal, while the y dimension was normalized for each quadrant using the outermost points of the white matter (Ronzano et al., 2022). The z-coordinates were normalized using the rostro-caudal positions of the rostral and caudal end of the mapped segments. The resulting cylindrical reconstruction of the spinal cord was then scaled to an idealized spinal cord size for the cervical, thoracic and lumbar part independently for illustrational purposes. All coordinate scaling and mapping were performed using a custom script in R adapted to read .csv files generated by Imaris (R Foundation for Statistical Computing, Vienna, Austria, 2005, http://www.r-project.org, version 3.6.2).

Analysis of axonal projections from lumbar dI3 interneurons in the cervical cord was performed using an adapted version of spinalJ (Fiederling et al., 2021). Briefly, a machine-learning pixel classification using Ilastik (version 1.3.3post3) was used to identify axonal processes. The probabilities of each pixel to be from a labelled axon or from the background were extracted from Ilastik. Pixels were classified as signal from axonal projection when the probability for them to be part of a labeled axon was higher than probability of background. To map the location of the projections to an annotated atlas of the spinal cord, the image registration to the reference atlas was then performed using Elastix (version 5.0.0) and a second channel imaged from neurotrace staining. The axonal projections were then mapped onto the Allen Brain Atlas Common Coordinate Framework. Visualizations of the data were performed in ImageJ, and subsequent analysis were performed in R.

The colocalisation of the GFP signal targeted to the presynaptic terminals with vGlut2 transporter was quantified manually on a total of 13 images (101.4x101.4µm) per animal, taken from a minimum of 5 different sections. For each animal, 5 images were taken from the region of dense dI3 ascending projections in C7 and C8 around motor neurons innervating the cutaneous maximus and 8 areas were scanned out of lamina 9 distributed randomly along C3 to T1. The number of presynaptic boutons apposed to retrogradely labelled motor neurons was quantified manually on a stack spanning the whole volume of the cells. The position of each motor neuron was recovered using the image of the entire hemisection acquired with a x10 air objective prior to the acquisition at high magnification, followed by the same pipeline used to generate distribution maps.

### Cutaneous maximus trunci reflex testing

Animals (1-2 months old) were first anesthetized with isoflurane and shaved. Two lines of dots were drawn on each side of the animal 0.5 and 1cm from the midline respectively and EMG probes were implanted when needed (see EMG recording section). Animals were then sedated using an intraperitoneal injection of pentobarbital at 30-35mg/kg (Dolethal 200mg/ml, vetoquinol), since the cutaneous maximus trunci reflex is resistant to pentobarbital (Petruska et al., 2014; Theriault & Diamond, 1988b), but abolished by ketamine/xylaxine, isoflurane and variable with urethane. When the sedation was wearing off, sedation was supplemented by additional intraperitoneal administration of 10mg/kg. Few minutes after pentobarbital administration and interruption of the isoflurane flow, reflexive skin twitches systematically recovered. This recovery was tested applying pinprick with a rigid 30G needle on the midline before starting to perform the tests with experimental stimuli. To assess cutaneous maximus trunci reflex, two Von Frey filament of 0.9g and 2g were used as well as two pinprick tests consisting of the tip of 30G needles glued to custom made Von Frey filaments of 0.9g and 2g (De Sousa et al., 2014). The stimuli were applied on one side of the animal, alternatively in the zone delineated by the midline and first line of dots and then between the two lines of dots. The stimuli were always applied in the posterior half of the animal. The sequence of the different stimulus was applied in series of each type of stimulus, with at least 15 seconds between each stimulus. Skin movements were recorded at 100 frames per seconds by a machine vision camera (FLIR, blackfly-S-BFS-U3-16s2M) using Spin View (Teledyne vision solutions, version 4.2.0.88) and Bonsai custom script (Bonsai Foundation, version 2.8.5, Lopes et al., 2015). The camera was placed on the side of the stimulation, and the contralateral responses were recorded using a mirror placed on the other side of the animal. The presence of skin twitches on each side of the animal was assessed by two independent raters, and the result for each trial was included only if both raters agreed on the outcome. For each type of stimulus, animals were included in the analysis of the percentage of responses only if they had a minimum of 10 trials with agreement between raters, then the percentage of response was calculated per animal and stimulus type.

The amplitude of the twitches was assessed for all trials with a skin twitch reflex using DeepLabCut (Mathis et al., 2018, version 3.0.0rc9) to track dots placed on the skin of the animals. Six dots per line were tracked (total of 24, 12 on each side) as well as landmarks to convert pixels in actual distance for the ipsilateral and the contralateral view. A ResNet-50-based was used with default parameters. Dot trajectories were then extracted using a custom R script allowing to extract peak of movements with a maximum amplitude. Movements were calculated defining individual axis for each line of dots, then maximum amplitude movements were detected when a minimum of 3 dots moved within a window of 8 frames by at least 5 times the standard deviation of their baseline (calculated from 30 successive frames before sensory stimulus). For each trial, the peak extracted was then manually verified to validate that the movements extracted corresponded to an actual response and was not due to a global movement of the animal (e.g. gasping) or an artefact of movements due to the stimulus itself moving the skin. After the peak detection was manually validated, the amplitude of the twitch on each line of dots was recorded as the dot with the highest amplitude of movement. When the movements of the dots were too small to be detected but a response had been recorded by manual assessment the values of the amplitude of movement was fixed to 0.

To determine the distance between the sensory stimulation and the center of the contraction, the rostro-caudal distance was manually measured (using ImageJ) between the point where the tip of the needle initially touched the skin and the location were the skin converged in opposite directions. To ensure that the small skin incision would not alter the localization of the motor response relative to the sensory stimulus, these experiments were performed on a cohort of animals in which no EMG probes were inserted. To make sure that the distance was not specific to one specific rostro-caudal level, the sensory stimulations were performed randomly along the rostro-caudal region covering the second third of the animal body measured from the neck to the base of the tail.

### EMG recording

A small incision of the skin was performed on each side on the posterior third of the animal 0.5 to 0.8cm from the midline. One myomatrix probe (RF-4x8-BVS-8, Camber) was then inserted under the skin bilaterally, rostrally to the incision along the rostro-caudal axis, while the ground was placed on the base of the tail. The myomatrix array was then connected to a RHD headstage (C3324, Intan technologies) coupled to a RHD 512-channel recording controller (C3004, Intan technologies) with a SPI interface cable (C3216, Intan technologies). To visualize the approximate timing of the stimulus on the EMG recording a manual trigger was sent to the recording controller when the stimulus was applied using a current stimulator (DS3, Digitimer). Recordings were performed using Intan RHX software (version 3.3.2) at 20kHz and band-pass filtered at 1-8000 Hz. Differential EMG signals were obtained by subtracting the raw signal from each neighboring channel from the raw signal of the channel of interest.

### Optogenetic

Optogenetic experiments were performed 4 to 5 weeks after intraspinal injections. The insertion of the cannula (CFMC22L10, Thorlabs) was made under general anaesthesia with isoflurane (5% for induction, 1-1.5% maintenance). Animals were placed in a custom device for stabilisation of the vertebrae, a laminectomy of C7 vertebrae was performed, the dura was removed and a small incision was made rostro-caudally at the point of cannula insertion to facilitate its insertion. The cannula connected to the optic fibre (M84L01, Thorlabs) using an interconnect (ADAF2, Thorlabs), was attached to a stereotaxic frame (Narishige, SMM-200) and subsequently inserted on the top of the ventral cord (750µm medio-laterally, 1000µm dorso-ventrally) in spinal segments C7 to C8. Once the cannula was in place the isoflurane anaesthesia was turned down and the mice were sedated using an intraperitoneal injection of pentobarbital at 5 mg/kg (Dolethal 200mg/ml, when the sedation was wearing off, sedation was supplemented by additional intraperitoneal administration). After the isoflurane anaesthesia had worn off, only red lights were kept on in the room and optogenetic stimulations were triggered using a 470nm LED (M470F1, Thorlabs) controlled with Spike 2 (version 9) and (CED Power 1401, Digitimer) with a minimum of 5 seconds between each pulse. While stimulations were applied, movements of the skin marked with a line of dots placed 0.5 cm from the midline and spaced by about 0.5 cm rostro-caudally, were recorded at 200 frames per seconds by a machine vision camera (FLIR, blackfly-S-BFS-U3-16s2M) controlled using Spin View (Teledyne vision solutions, version 4.2.0.88) and Bonsai custom script (Bonsai Foundation, version 2.8.5). This procedure was performed similarly in Isl1^cre/wt^ and Cre negative animals. In Cre negative animals, the cannula was placed successively in 3 different sites rostro-caudally to make sure that the absence of response was not due to a misplacement of the stimulation.

The amplitude of skin twitches was analysed using DeepLabCut (version 3.0.0rc9) using a method similar to the one described for the cutaneous maximus reflex testing. Only one line of dots was tracked on the side ipsilateral to the optogenetic stimulation with skin movements measured along the axis defined by the line of dots. In addition, light pulses were tracked to define the time window in which skin twitches were measured. The baseline was calculated from a window of 45 frames preceding light pulse, and a movement was considered genuine when one of the dots moved by at least 5 times the baseline standard deviation. For each trial, the peak extracted was then manually verified to validate that the movements extracted corresponded to an actual response and was not due to a global movement of the animal (e.g. gasping). The presence of skin twitches upon optogenetic stimulation was assessed by two independent raters, and the result for each trial was included only if both raters agreed on the outcome. For each type of pulse length, animals were included in the analysis of rate of response only if they had a minimum of 5 trials with agreement between raters, then the response rate was calculated per animal and pulse length.

### Statistical analysis

Statistical analyses were performed using R and R Studio (v1.4.1717). The difference between groups were considered significant for *p* < 0.05. The statistical tests used are define in the figure legend and the level of significance are reported as follow: \**p* < 0.05, \*\**p* < 0.01, \*\*\**p* < 0.001, \*\*\*\**p* < 0.0001, ns. indicates no significance. In the text, summary data are reported as mean ± SD.

Most of the data produced using behavioural or anatomical methods have an intrinsically nested structure, in which observations are taken from the terminals/cells/trials, in a specific animal. Therefore, to analyse the data collected, the origin of the variance in our datasets was identified. The interclass correlation coefficient (ICC) was calculated using a one-way random effect ICC model (1,1) with the ICCbare function of the ICC R package. When ICC≤0.5 in all the groups to be compared, data were treated as independent and applied all the following tests to the lowest level of observation. When ICC≥0.5 in at least one of the group, a hierarchical bootstrap was performed. First animals (first level) were resampled with replacement and then within each animal the terminals/trials were resampled with replacement (second level). After ICC calculation, the normality of the distributions was tested using a Shapiro-Wilk tests and the equality of variance using a Levene test to select adapted statistical tests. When n<8 non-parametric tests were used.

Bootstrap replicas were computed with paired or non-paired resampling depending on the structure of the data and the Hedges’s G coefficient calculated using the R function “cohens.d” from the “effsize” package. For behavioural data, when values were plotted per animal, bootstrapped resampling was restricted to the computation of Hedge’s G coefficient. The Hedge’s G reported correspond to the median of 5000 bootstrap replicas (with a sample of maximum 1000 replicas per group), that were performed with a resampling with replacement. All Hedge’s G coefficient are reported as absolute values. When bootstrapping was used for computing p values, the reported p values correspond to the median of the distribution.

---

**Supplementary figure 1:**
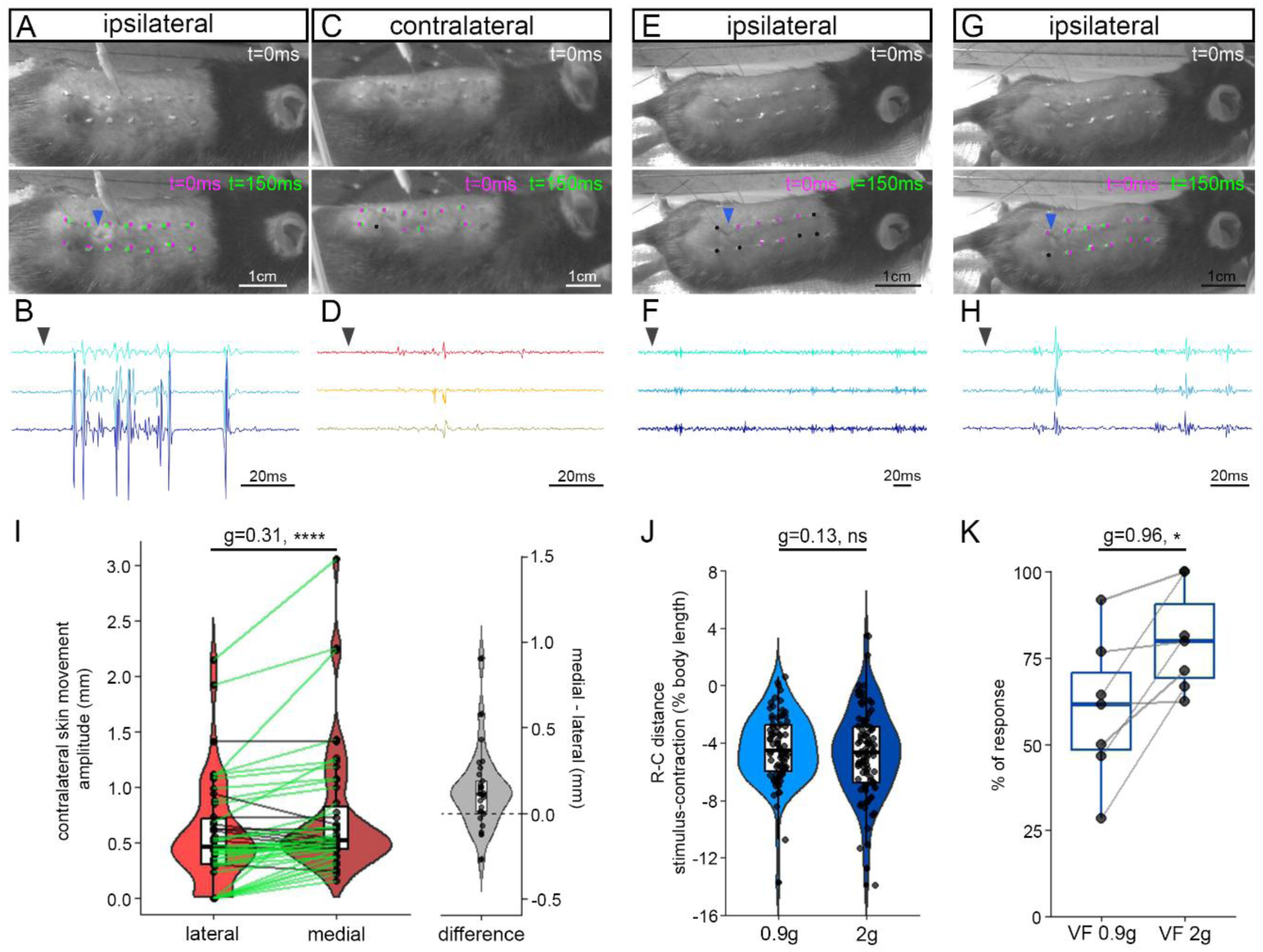
reflexive skin twitches rely on a spatial representation of the torso. (A, C, E, G) Example of recorded skin movements following stimulation with a 2g pinprick on the side ipsilateral (A) and contralateral (C) to the sensory stimulation. Example of recorded ipsilateral skin movements following stimulation with 0.9g (E) and 2g (G) Von Frey filaments. Images at t=0ms are from the first frame with contact of the pin/Von Frey on the skin, while 150ms later the skin shows a twitch. Magenta and green dots highlight the position of dots placed on the skin to visualize the twitch (magenta for t=0ms, green for t=150ms). Black dots indicate no movement between the two time points. The blue arrowhead points at the position of initial muscle contraction. (B, D, F, H) Differential myomatrix EMG signal recorded from cutaneous maximus from responses showed in the panels above. The black arrowhead indicates the approximate timing of sensory stimulation. (I) Maximum amplitude of skin movement on the side contralateral to the pinprick, on the lines of dots drawn on the lateral versus medial part of the cutaneous maximus (left). Difference of amplitude of contralateral skin movement between the medial and lateral dotted lines (right). Individual trials are connected by green lines when showing a higher value on the medial part of the cutaneous maximus while black lines are used to join trials with higher amplitude on the lateral part of the muscle. Trials from 0.9g and 2g pinprick stimulations are pooled together. (J) Rostro-caudal distance expressed in percent of body length between the sensory stimulation and the location of the initial contraction of the cutaneous maximus following stimulation with a 0.9g or 2g pinprick. When contraction is caudal to the sensory stimulus the percentage is set as negative, and positive when the shift is rostral. (K) Percentage of trials leading to an ipsilateral skin twitch following stimulation with 2g and 0.9g Von Frey filaments. (I) n = 47 responses (18 from 0.9g and 29 from 2g) in a total of N=8 mice. Non-hierarchically bootstrapped paired Hedge’s G coefficients and two-tailed paired Wilcoxon test. (J) n = 92 (2g), and 80 (0.9g) responses in a total of N=5 mice. Non-hierarchically bootstrapped Hedge’s G coefficients and two-tailed Mann-Whitney test. (K) Individual values plotted from n=7 mice, two-tailed paired Wilcoxon test and paired bootstrapped Hedge’s G coefficients.

**Supplementary figure 2:**
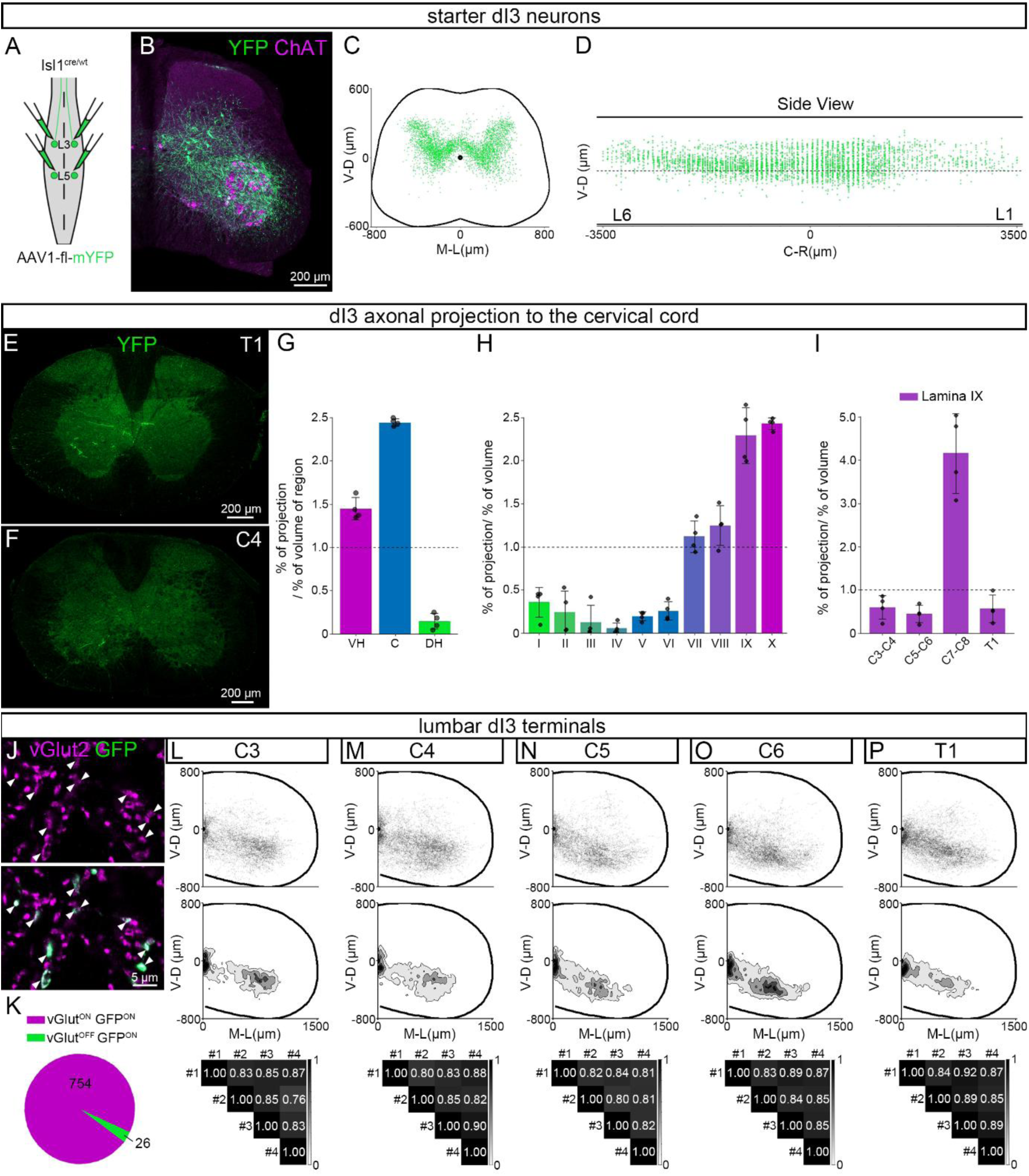
lumbar dI3 neurons form long ascending projections to cutaneous maximus motor neurons. (A) Experimental strategy used to map axonal projections of lumbar dI3 neurons in cervical segments of the spinal cord. (B) Orthogonal projection showing somata of transduced lumbar dI3 neurons (YFP, green) and cholinergic neurons (ChAT, magenta) on a lumbar cord transverse section. (C-D) Spatial distribution of YFP^ON^ChAT^OFF^ lumbar dI3 neurons on an idealized lumbar transverse section (C) and sagittal section (D, both sides of the cord overlapped). (E-F) Orthogonal projections showing lumbar dI3 axons (YFP, green) on transverse sections from cervical segment C4 (E) and thoracic segment T1 (F). (G) Percentage of lumbar dI3 neurons axonal projections scaled by the volume of each region in cervical segments C3 to T1, between ventral horn (VH), central canal area (C) and dorsal horn (DH). (H) Percentage of lumbar dI3 axonal projections in cervical segments (C3 to T1) across spinal cord laminae scaled by their volume. (I) Distribution of lumbar dI3 axonal projections along lamina IX across segments C3 to T1 scaled by its volume. (J) Single optic slice from lumbar dI3 terminals (GFP, green) in lamina IX of caudal cervical segments showing their expression of vGlut2 (magenta). (K) Distribution of lumbar dI3 neurons terminals (GFP^ON^) colocalized (magenta) or not (green) with vGlut2 signal. (L-P) Spatial distribution, spatial density and spatial correlation analysis of lumbar dI3 terminals in segments C3 (L), C4 (M), C5 (N), C6 (O) and T1 (P) on idealized cervical transverse sections. (C-D, L-P) Spatial distributions and densities are pooled from tracing in n=4 mice. (G-I) Data shown are from n=4 mice, with individual values plotted. (K) Numbers on the pie chart correspond to individual terminals quantified in a total of n=5 mice.

**Supplementary figure 3:**
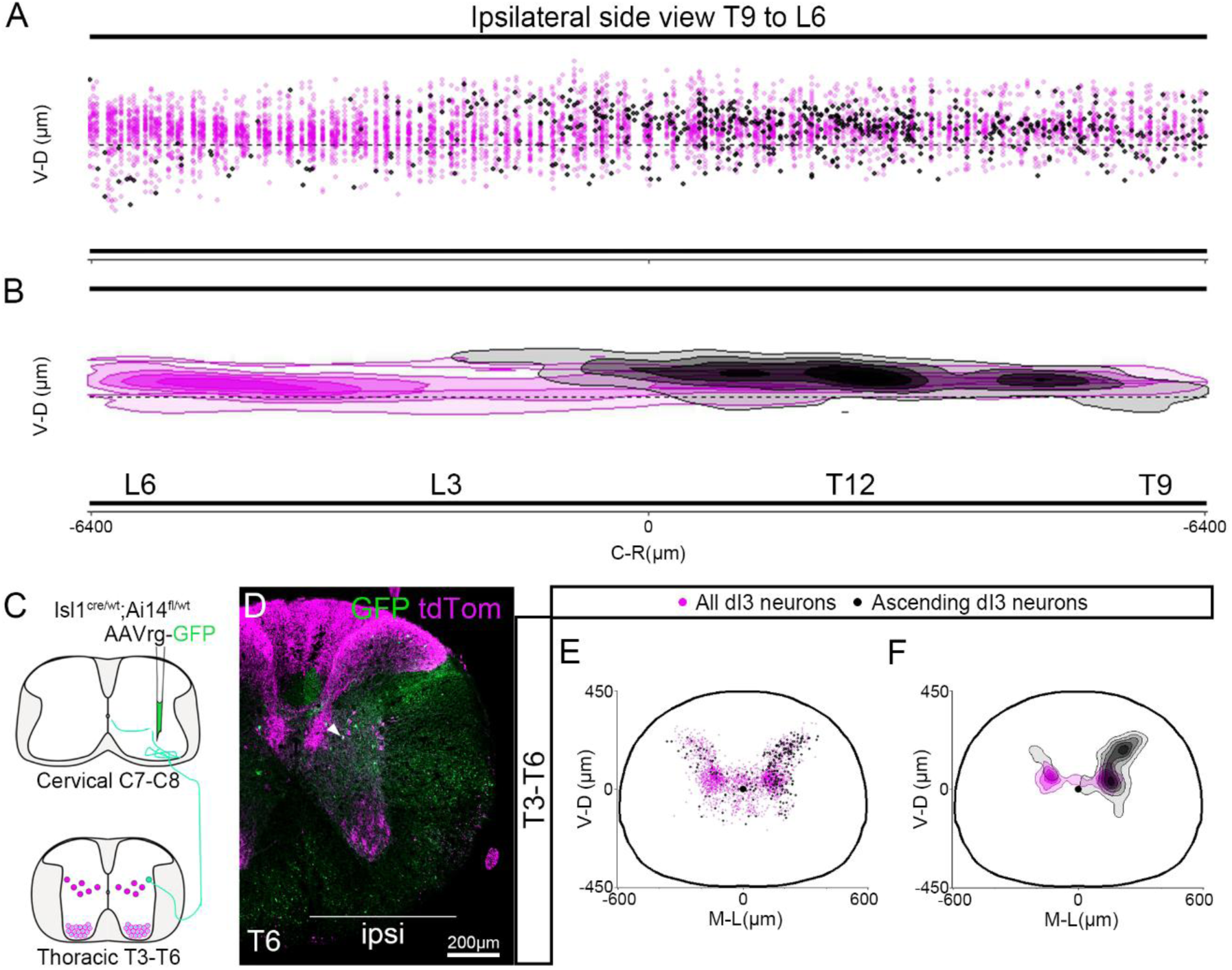
Long ascending projections are formed by a subset of dorso-lateral dI3 neurons. (A-B) Spatial distribution (A) and density (B) of all ipsilateral dI3 neurons (magenta) and ipsilateral dI3 neurons projecting to the ventral horn of segment C7-C8 (black) on an idealized sagittal section from segments T9 to L6. (C) Experimental strategy used to map on rostral thoracic segments T3 to T6 the global distribution of dI3 neurons (tdTomato^ON^) and the ones projecting to the ventral horn of C7-C8 segments (GFP^ON^tdTomato^ON^). (D) Orthogonal projection of ipsilateral hemislices showing a GFP^ON^tdTomato^ON^ dI3 neuron (GFP in green, tdTomato in magenta) from thoracic segment T6. (E-F) Spatial distribution (E) and spatial density (F) of all dI3 neurons (magenta) and dI3 neurons projecting to the ventral horn of segment C7-C8 (black) from segments T3 to T6 on an idealized transverse thoracic section. Results pooled from n=4 mice, mapped on every other sections for GFP^ON^tdTomato^ON^ dI3 neurons while the global distribution of dI3 neurons (tdTomato^ON^) was mapped on one section every four.

**Supplementary figure 4:**
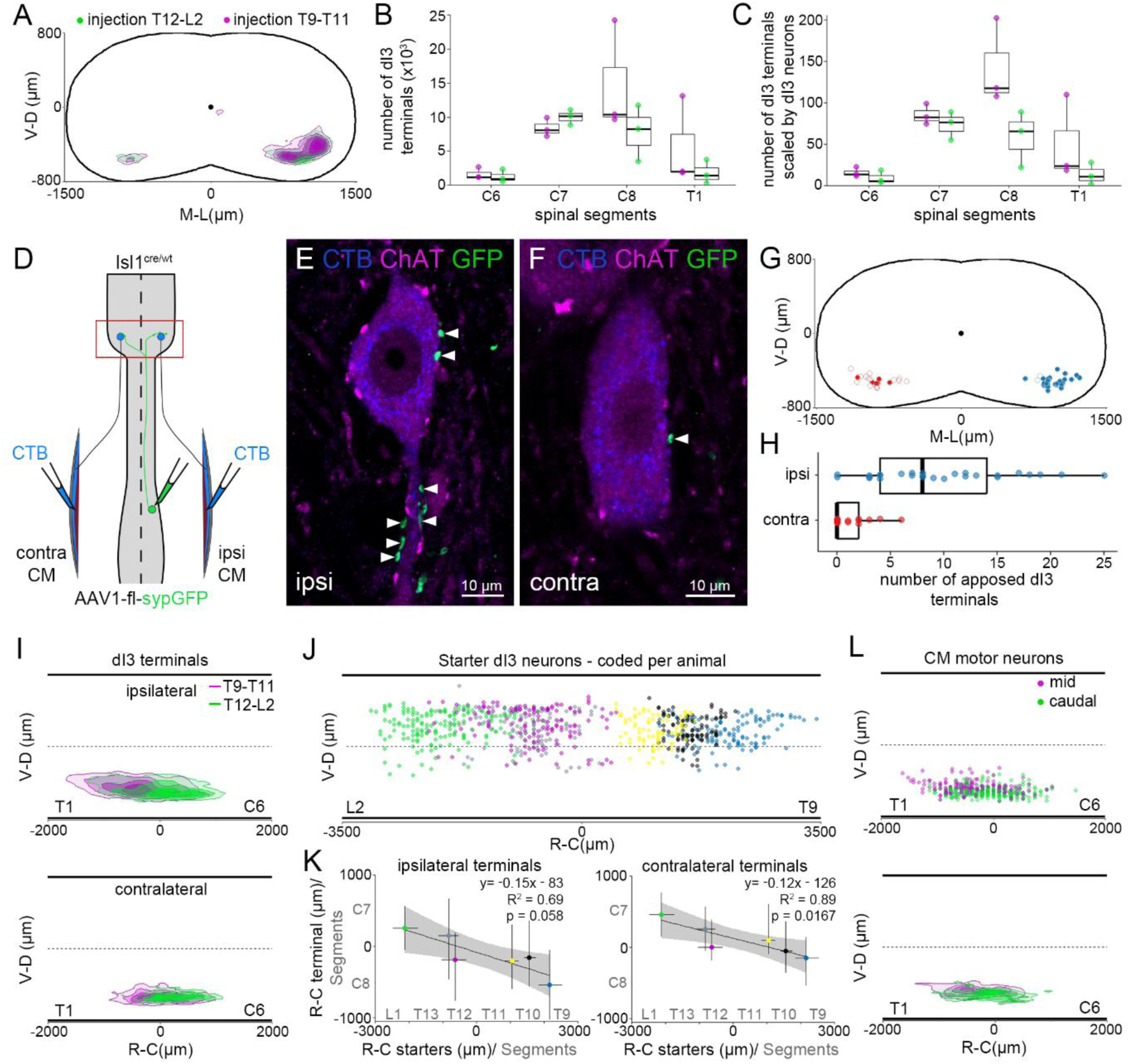
Cutaneous maximus motor neurons and ascending dI3 projections are somatotopically organized along the rostro-caudal axis. (A) Spatial density of GFP presynaptic terminals from dI3 neurons in T12 to L2 (green) and T9 to T11 (magenta) mapped from segments C6 to T1 on an idealized cervical transverse section. (B-C) Number of dI3 neurons terminals (in thousands, B) and number of dI3 terminals scaled by GFP positive dI3 neurons (C) quantified in every other sections following tracing of dI3 neurons in T9 to T11 (magenta) and T12 to L2 (green). (D) Experimental strategy used to visualize presynaptic terminals from lumbar dI3 neurons on ipsilateral and contralateral identified cutaneous maximus motor neurons. (E-F) Single optic slice showing presynaptic boutons (GFP, green) from dI3 neurons located in segments T12 to L2 on an ipsilateral (E) and a contralateral (F) motor neuron (ChAT, magenta) innervating the cutaneous maximus (CTB, blue). White arrowheads highlight dI3 boutons apposed to the cutaneous maximus motor neurons. (G) Position of analysed ipsilateral (blue) and contralateral (red) cutaneous maximus motor neurons on an idealized cervical transverse section. Filled circles are motor neurons with apposed terminals from dI3 neurons of segments T12 to L2, while unfilled circles correspond to motor neurons with no apposed terminals. (H) Number of apposed terminals from dI3 neurons in segments T12 to L2 on ipsilateral vs contralateral cutaneous maximus motor neurons. (I) Spatial density of presynaptic terminals from dI3 neurons of T9 to T11 segments (magenta) and T12 to L2 segments (green) along the rostro-caudal axis on an idealized sagittal section from segments C6 to T1. Densities are plotted independently for terminals on the ipsilateral (top) and contralateral (bottom) side to the injection. (J) Spatial distribution of dI3 neurons transduced by AAV1-fl-sypGFP injections, plotted on an idealized sagittal section from T9 to L2 and colour coded per animal. (K) Correlation per animal between the median of the rostro-caudal distribution of transduced dI3 neurons somata in thoraco-lumbar segments and the median of the rostro-caudal position of their synaptic terminals mapped from segments C6 to T1, on the ipsilateral (left) and contralateral (right) side of the cord. Animals are colour coded as in panel J. (L) Spatial distribution (top) and density (bottom) of motor neurons innervating the caudal (green) and middle (magenta) part of the cutaneous maximus along the rostro-caudal axis on an idealized sagittal section from segments C6 to T1. (A) Spatial densities are pooled from tracing in n=3 mice per type of injection. (B-C) Data shown are from n=3 mice per type of injection. (G-H) n= 27 ipsilateral and n=21 contralateral cutaneous maximus motor neurons from N=3 mice. (I, L) Spatial distributions and densities are pooled from tracing in n=3 mice for dI3 terminals and n=4 mice for cutaneous maximus motor neurons. (K) R^2^ and p values were calculated using the Spearman rank correlation coefficient.

***Supplementary video 1: reflexive skin twitch following stimulation with a 0.9g pinprick*.**

Video recorded at 100 frames per seconds, speed is slowed down at 0.2x.

***Supplementary video 2: reflexive skin twitch following stimulation with a 2g pinprick*.**

Video recorded at 100 frames per seconds, speed is slowed down at 0.2x. ***Supplementary video 3: reflexive skin twitch following stimulation with a 0.9g Von Frey filament.*** Video recorded at 100 frames per seconds, speed is slowed down at 0.2x.

*Supplementary video 4: reflexive skin twitch following stimulation with a 2g Von Frey filament.* Video recorded at 100 frames per seconds, speed is slowed down at 0.2x.

***Supplementary video 5: Skin twitch following optogenetic stimulation of dI3 neurons ascending projections.*** Video recorded at 200 frames per seconds, speed is slowed down at 0.2x. Optogenetic stimulation with a single pulse of 20ms.

